# Flat-field super-resolution structured illumination microscopy with joint spatial-temporal light modulation

**DOI:** 10.1101/2024.05.01.591776

**Authors:** Yong Liang, Gang Wen, Jie Zhang, Simin Li, Yao Tan, Xin Jin, Linbo Wang, Xiaohu Chen, Jing Gao, Hui Li

**Affiliations:** Jiangsu Key Laboratory of Medical Optics, Suzhou Institute of Biomedical Engineering and Technology, Chinese Academy of Sciences, Suzhou, Jiangsu, 215163, China; Cixi Institute of Biomedical Engineering, Ningbo Institute of Materials Technology and Engineering, Chinese Academy of Sciences, Ningbo 315201, PR China; College of Chemistry, Chemical Engineering, and Materials Science, Shandong Normal University, Jinan 250014, China

## Abstract

Super-resolution structured illumination microscope (SR-SIM) has been established as a powerful tool for visualizing subcellular dynamics and studying organelle interactions in live cells. However, the interfering Gaussian beams result in a limited and nonuniform field of view (FOV) which hinders its application for large whole-cell dynamics and pathological sample imaging. Here, we proposed a joint spatial-temporal light modulation (JSTLM) method to reshape the excitation light field into flat-field structured illumination without disturbing the interfering fringes. Our flat-field structured illumination microscopy (flat-field SIM) improves the uniformity across the whole FOV significantly, hence enabling SR image stitching. Skeleton dynamics and vesicle transportation in and between whole cells were visualized by flat-field SIM. With the stitching of multi-FOV flat-field SIM images, millimeter-sized SR images can be obtained which provides the possibility for cell heterogeneity studies and pathological diagnoses. The JSTLM method can be further incorporated with regions of interest to reduce unnecessary photodamage to live cells during multicolor imaging.

**Contributions:** Y.L. and X.H.C. conceived and designed the idea. Y.L., S.M.L., X.J., and G.W. built the SIM setup. Y.L. performed the data acquisitions. Y.L. and X.H.C. conducted the optical wave simulation. J.Z. prepared the cell samples. Y.T. and L.B.W. performed the image analyses. Y.L. prepared the illustrations. X.H.C. and J.G. supervised the project. Y.L. and H.L. wrote the manuscript.

## Introduction

In the past 20 years, super-resolution fluorescence microscopy (SRFM) has renewed the optical imaging field by breaking the diffraction limit and led to significant advances in biological research^1–6^. Among the various super-resolution techniques, such as single-molecule localization microscopy (SMLM)^2,3^, stimulated emission depletion (STED)^4^, expansion microscope^5^, structured illumination microscopy (SIM)^6^ hold special advantages in the aspect of imaging speed, photon efficiency, and compatibility with generally-used labeling strategies. With the rapid improvement in the SR-SIM instrumentation and image reconstruction algorithms, it has been well acknowledged by biological society to observe the subcellular structures, monitor the cellular dynamics, and discover new regulation mechanisms^7–10^. Despite the existence of various SIMs, such as Fair-SIM^7^, Open-SIM^8^, Hessian-SIM^9^, GI-SIM^10^, HiFi-SIM^11^, and PCA-SIM^12^, the underlying principle for the generation of structure illumination primarily relies on the interference of two or three Gaussian laser beams (referred to as Gaussian-SIM in this paper)^6,13^.

Beyond the observation of individual or a few organelle structures and dynamics, their structure variance and distribution in a whole cell or among multiple cells are also important for cell biology studies^14,15^ and especially for pathological diagnosis. The extension of SRFM to high-throughput and high-content imaging triggers new research in a wide range of biomedical areas^16,17^, such as cell fate determination, drug treatments, and pathological investigation. In these researches, imaging with a large field of view (FOV) even at a millimeter scale for hundreds of cells is highly demanding.

The illumination uniformity in the FOV is critical for large-field imaging and further stitching. However, the lasers that are used in SRFM as excitation light sources mostly are Gaussian beams. It results in uneven illumination at the focal plane with the intensity profile following a Gaussian distribution. The uneven illumination leads to position-dependent resolution, limits the effective FOV, and hinders the quantitative analyses based on the fluorescence intensity^18–26^. It also causes dimmed borders surrounding each image during stitching and poses significant challenges for the image mosaic algorithm^26^. So many studies have been conducted to develop flat-field illumination for SRFM^18–25^. Beam-shaping devices^18,19^, such as a Pishaper or Topshape, can directly transform the Gaussian distribution to a flat-field beam. Optical waveguides^20,21^ can provide uniform illumination across large FOVs but have fixed illumination depths and sizes. Square core fibers^22^ and microlens arrays^23,24^ can also provide efficient homogeneous illumination over large FOVs. ASTER^25^, which scans a Gaussian beam at a fixed gap by using 2D galvanometers to generate flat-field illumination in a time-averaged manner, is an effective approach to producing uniform excitation. The above techniques have been successfully applied to epifluorescence microscopy, SMLM^18–23,25^, and multifocal structured illumination microscopy^24^.

The Gaussian-shaped beams also hold a similar limitation for SR-SIM and the situation is even worse. In current Gaussian-SIM, spatial light modulators (SLM)^27,28^ are usually used to generate two or three interference beams, so FOV is constrained by not only the size of the SLM but also the profile of the input Gaussian beam. Nonuniform Gaussian illumination leads to a decrease in signal-to-noise ratios (SNR) at the edge of FOV compared to its central region. Reconstruction artifacts in SIM occur frequently with low SNR so the quality of the reconstructed image at edge is greatly reduced^9,11^. In practice, only the central portion of the Gaussian beam is utilized to mitigate the position-dependent reconstruction quality. Consequently, the FOV is significantly reduced compared to that in SMLM. Moreover, the above methods for flat-field illumination^18–25^ cannot be directly applied to SIM because the wavefront will be altered in these flat-field modulation methods, causing distorted interference patterns. To our knowledge, by now there is no successive method for achieving high-quality flat-field structured illumination that has been reported for laser interferometric SIM.

In this paper, we developed a joint spatial-temporal light modulation (JSTLM) technique to modulate the laser illumination intensity distribution in the FOV while maintaining the plane wave wavefront of the laser beams. By decomposing the modulation function of the flat-field illumination into a set of binary bit planes with different time weights, the Gaussian-shaped interference pattern was transformed into the flat-field fringe pattern. So an effective FOV of four times larger was demonstrated, with the possibility to be further increased with the limitation only by the size of SLM. With this flat-field SR-SIM, we tracked the dynamics of hundreds of vesicles in a live whole cell. The flat-field SR-SIM also enables image stitching to perform SR imaging at a millimeter scale, which covers hundreds of cells to study their mitochondria structure heterogeneity and to image pathological slide samples. Furthermore, the JSTLM technique can enable selective flat-field SIM at user-defined regions of interest (ROI). We demonstrate two-color SR-SIM imaging for one laser illuminating the cell nucleus region and the other laser illuminating the cytoplasmic region, which photodamage to live cells is greatly reduced by this selective SIM technique.

## Results

### Principle of joint spatial-temporal light modulation

In SR-SIM, the excitation light intensity in the focal plane can be expressed as

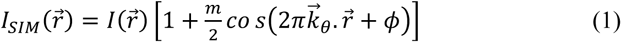

where 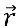 is the spatial coordinate; 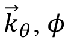 and *m* are the spatial frequency, the initial phase, and the modulation depth, respectively. Generally, the distribution of *I* 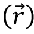 was neglected and set as a global constant *I_0_* at the entire FOV. However, in reality, 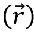 has a distribution according to the excitation beam shape. In the Gaussian-SIM, it was displayed as a Gaussian-shaped distribution.

To achieve high imaging speed, recently developed SR-SIM usually uses a ferroelectric liquid crystal SLM (FLC-SLM) to diffract the input Gaussian beam into two or three coherent beams, as depicted in Fig. 1a. The FLC-SLM employs only 0 and 𝜋 phases and has a switching speed on the order of microsecond, much fast than the sCMOS camera. In 2D-SIM imaging, the light field is modulated by preloading nine hologram bit plane images to FLC-SLM with different orientations and phases. As the loaded bit plane is binarized, the illumination intensity exhibits a Gaussian-shaped distribution the same as the input beam (Fig.1b). To modulate the illumination intensity distribution, we proposed a joint spatial-temporal light modulation method. This method modulates the effective illumination intensity distribution by loading a set of bit planes with different time weights within a one-frame camera exposure time; the details are presented in Supplementary Note 1. Hence, the illumination pattern in the flat-field SIM can be denoted as

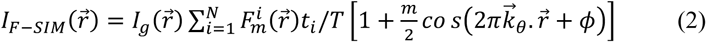

where 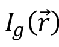 represents the Gaussian-shaped incident light; 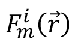 denotes the *i_th_* bit plane of the modulation function; *t_i_*is the illumination time weight of the *ith* bit plane 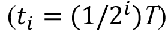; *T* is the exposure time of each frame; and *N* represents the total number of bit planes.

**Fig. 1.**
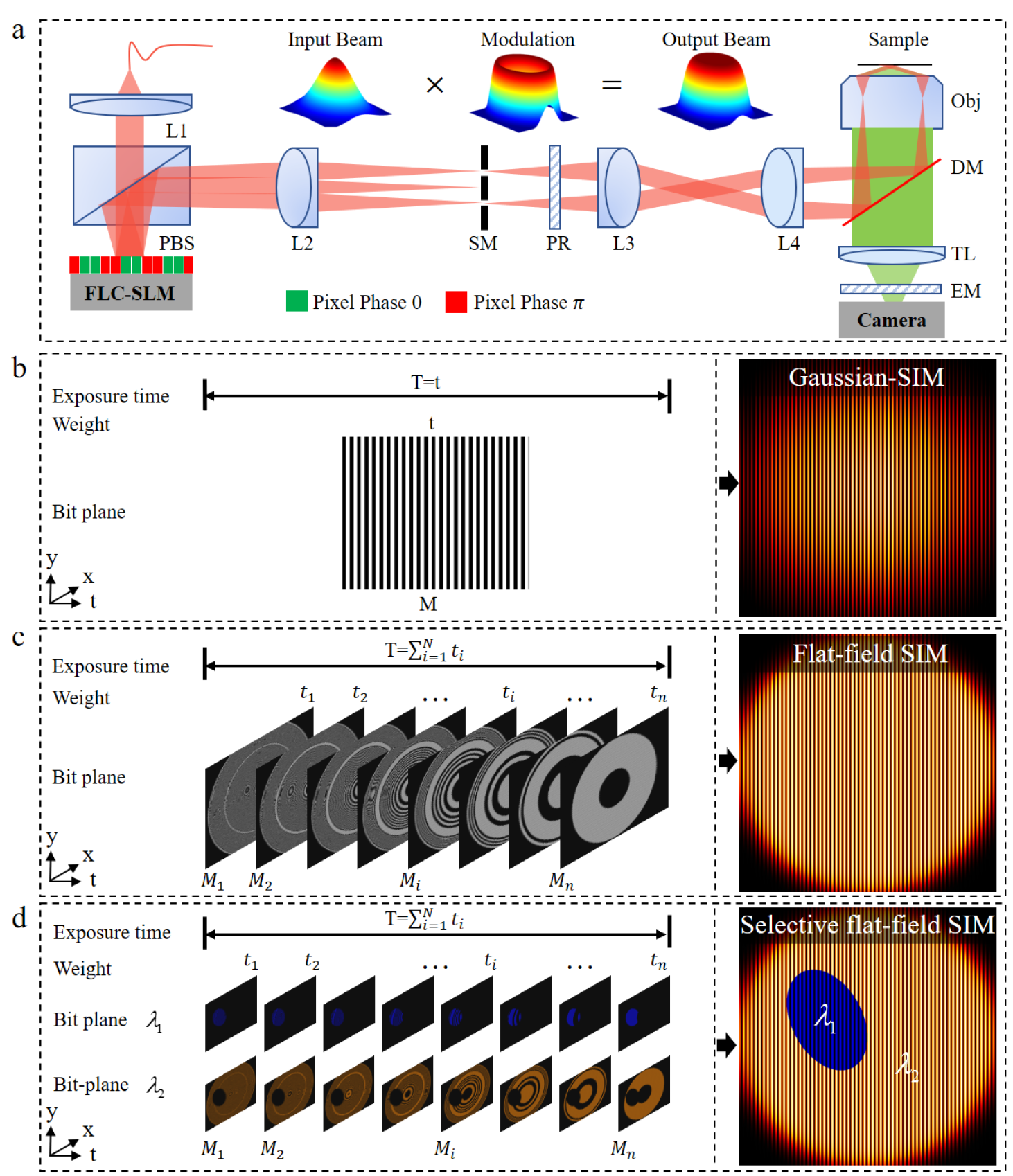
Principle of the joint spatial-temporal light modulation method for flat-field SIM. a. Flat-field SIM is implemented in a classic SIM system based on a binary FLC- SLM. A modulation function is applied to the input Gaussian beam to transform it into an effective flat-field intensity distribution. **b** Binary bit plane loaded on the FLC-SLM to generate fringe patterns in the Gaussian-SIM. **c** Sequence of bit planes with varying illumination weights loaded on the FLC-SLM to generate flat-field fringe patterns in the flat-field SIM. **d** Two sets of bit plane sequences loaded on the FLC-SLM to generate selective flat-field fringe patterns at different regions of interest for two-color imaging.

Assuming a Super-Gaussian function with the order of 8 as an approximation of flat-field distribution, the intensity modulation function 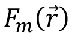 can be obtained explicitly and then divided into a set of bit plane modulation function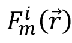 corresponding to different time weight *t_i_*. N bit-planes with fringes were generated for flat-field SIM, preloaded into FLC-SLM, and switched sequentially within one exposure time of the sCMOS camera (in the order of 10 ms) (Fig. 1c, Fig. S2). With this JSTLM method, the Gaussian beam can be modulated into an effective flat-field illumination with cosine-interference fringes. Furthermore, a user-defined region can be superimposed in the flat-field illumination to achieve selective flat-field SIM at regions of interest (Fig. 1d).

To verify the preservation of the interference fringe during the modulation of the Gaussian beam into a flat-field illumination, we conducted a wave optics simulation based on angular spectrum diffraction. Details about the simulation process and results are presented in Supplementary Notes 1.3 and 1.4. For each bit plane, the intensity at the focal plane shows cosine fringe distribution with high modulation depth (Fig. S3a), indicating that the JSTLM method maintains the wavefront of the incident plane wave during the modulation process. Importantly, the fringe patterns corresponding to 8 bit planes have the same period and phase so that they can be added together without decreasing the modulation depth. The final intensity distribution is obtained by a weighted summation of the illumination intensity of each bit plane, as shown in Fig. S3c. Compared to Gaussian illumination (Fig. S3b), the JSTLM enables an even illumination intensity over the entire illumination region while preserving the interference fringes with high modulation depth (Fig. S3d).

It should be pointed out that the uniformity of the flat-field illumination is correlated with the size of the pinholes, which are placed at the focal plane of the Fourier lens as spatial filters. In 2D Gaussian-SIM, the pinholes filtered out the high-order diffractions and retained only *±* 1-order beam. In flat-field SIM, the high-frequency components corresponding to the bit planes were also filtered out. To obtain the optimal illumination pattern, we conducted simulations using 74 pinholes of different sizes and established a quantitative correlation between the uniformity of the FOV and the pinhole size (Fig. S3e). The results show the optimal uniformity for flat-field structured illumination could be obtained with the pinhole size within 0.65%*K_max_* to 0.75%*K_max_*, where *K_max_* is the corresponding spatial frequency of the camera pixel size at the SIM mask plane.

To experimentally characterize the illumination uniformity, we imaged a thin layer of rhodamine fluorescent dye (details in Supplementary Note 2). Fig. S4 demonstrates that flat-field excitation induces more homogeneous fluorescence across a large FOV with interference fringes (shown in the subfigure). The fringe looks weak because its period was close to the diffraction limit and was convoluted by the points spread function during imaging. The coefficient of variation (COV, defined as the standard deviation divided by the mean) was calculated to quantitatively evaluate the uniformity of the illuminated light field^18^. The COV values for the flat-field and Gaussian distributions are 11.9% and 33.1%, respectively, indicating a three-fold improvement in the homogeneity. At 75% illumination uniformity, the flat-field illumination provides an FOV that is approximately four times larger than the FOV obtained by the Gaussian illumination.

## Performance of the flat-field SIM

To quantitatively assess the super-resolution imaging performance of flat-field SIM, fluorescent microspheres with diameters of 100 nm were imaged. As shown in Fig.2a, the beads closer to the center of the FOV had stronger fluorescence than those near the edge. In contrast, the fluorescence intensity of the beads exhibits a homogeneous distribution under flat-field illumination (Fig. 2b). Fig.2c and Fig.2d show the zoomed-in image at the center and edge, respectively. The intensity profile along the x-axis direction in Fig. 2e clearly shows that the intensity gradually decreases from center to edge for Gaussian-SIM while keeping roughly constant over a large region for flat-field SIM. The coefficient of variation, defined as the standard deviation divided by the mean, was 80% and 52% for the Gaussian and flat-field excitation. The difference in light intensity distribution can also be inferred from the photobleaching after 15 min imaging under Gaussian and flat-field illumination (Fig. S7). The rate of photobleaching was uniform with flat-field illumination, whereas the rate of photobleaching was faster in the center than near the periphery with Gaussian illumination.

**Fig. 2.**
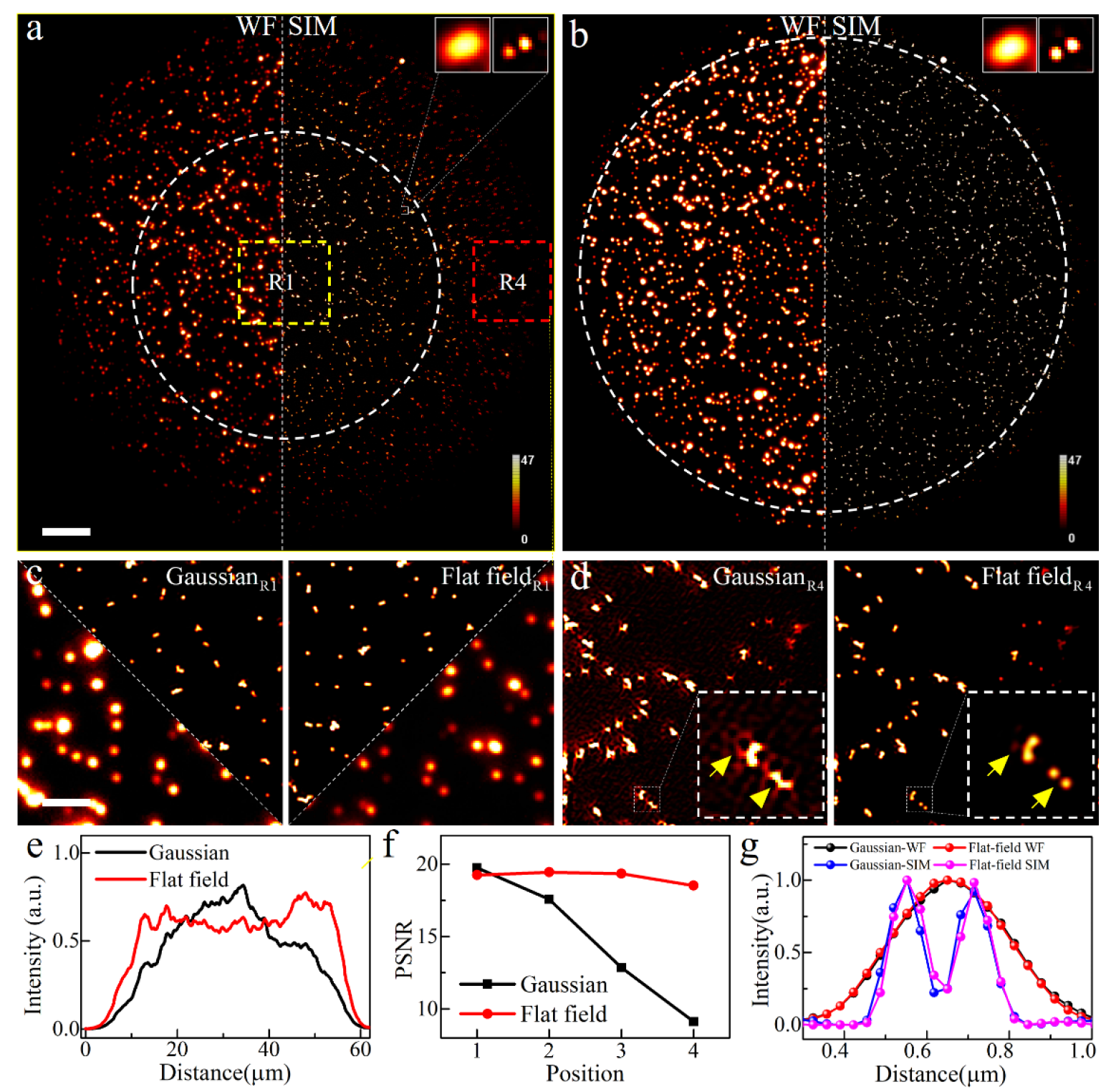
Super-resolution imaging of fluorescent beads with Gaussian-SIM and flat- field SIM. a. Image acquired by the Gaussian-SIM has a higher intensity at the center than at the edge for both an equivalent WF (left) and SR-SIM image (right). **B** Image acquired at the same position with the flat-field SIM shows a uniform fluorescence intensity across the FOV (indicated by the dashed circle) for both an equivalent WF (left) and SR-SIM image (right). **c** Magnified images show doubled resolution enhancement for both Gaussian and flat-field SIM images at the center of the FOV (R1, the yellow box in (a)). **d** Magnified images show reduced SR image quality with many hammerstroke artifacts for Gaussian-SIM and maintained high-quality SR image for flat-field SIM at the edge of the FOV (R4, red box in (a)). **e** Average intensity profile along the vertical direction for Gaussian and flat-field illumination. **f** PSNR of four subregions from the center to the edge (R1-R4). **g** Normalized intensity profile along two adjacent beads in the upper right corner of (a) and (b). Scale bar: 5 *μ*m in (a) and (b), 1.5 *μ*m in (c) and (d).

The even illumination also guarantees an equal signal-to-noise ratio (SNR) at the entire FOV, which is critical for super-resolution image reconstruction. It is well-known that SIM reconstruction is an ill-posed inverse problem that is highly susceptible to artifacts, particularly for raw data at low SNR^9–11,28,29^. For quantitative comparison, we localized and analyzed the peak SNR (PSNR) of each fluorescent bead under Gaussian and flat-field illumination with equivalent peak power. The average PSNR for all fluorescent beads under Gaussian illumination was 13.09, whereas this value significantly improved to 29.13 under flat-field illumination, representing a 2.2-fold enhancement. Four sub-FOVs spanning from the center of the FOV to the edge were selected to conduct further evaluation. Fig. 2f shows that the PSNR in the center FOV is 20.8 under Gaussian illumination; however, this value rapidly decreases with increasing distance from the center to 8.5 at the edge of the FOV, representing an approximately 60% decrease. However, the attenuation of the PSNR is only 10%, decreasing from 20.2 to 18.2 under flat-field illumination.

Due to decreased illumination intensity and SNR at the edge, the reconstructed SR image quality will be greatly impaired with Gaussian-SIM. Fig.2c and 2d show zoomed sub-FOVs at the center and edge for comparison. All beads are clearly resolved for flat-field SIM image (Fig. 2c) with a FWHM of 101*±*2.8 nm (Fig.2g). However, the image collected by the Gaussian-SIM exhibits significant noise-related artifacts (yellow arrow in Fig. 2d) at the edge of the FOV due to the low SNR (Fig. 2f). These hammerstroke artifacts^9,11^ occur frequently and hard to be eliminated reliably. So in practical application, these regions with strong artifacts were cut out and only SR images within a certain diameter were displayed. This operation results in a small FOV that hinders whole-cell dynamic imaging and a waster of laser dose that may introduce photodamage to live cells. In contrast, the even illumination intensity and SNR with flat-field SIM ensure the same SR image quality with an effective FOV four times larger, without change of any optics.

### Super-resolution imaging of cell skeleton by Flat-field SIM

To explore biological specimen imaging, the microfilaments in live U2OS cells were imaged with the Gaussian-SIM and flat-field SIM. As depicted in Fig. 3a, b, the flat-field SIM exhibits high-quality reconstruction comparable to that of the Gaussian-SIM in the center of the FOV. The magnified images (Fig. 3c) and normalized intensity profiles (Fig. 3e) show that microfilaments with a spacing of 124 nm can be distinguished in both the Gaussian-SIM and flat-field SIM images. However, the reconstruction results differ significantly at the edge of the FOV. Compared to the flat-field SIM images, the fine structure of the microfilaments cannot be resolved in the Gaussian-SIM images, and the images contain obvious residual noise-related hammerstroke artifacts (Fig. 3d). These differences are attributed to the low SNR at the edge of the FOV caused by the low illumination intensity. The absolute intensity profile confirms that flat-field SIM facilitates quantitative analysis based on fluorescence intensity. As shown in Fig. 3f, the difference in absolute fluorescence intensity between the peak and trough is 83 (33% of the peak intensity) in the flat-field SIM image, while this difference is only 13 (5% of the peak intensity) and nearly lost in local noise in the Gaussian-SIM image. The resolution was also calculated with the decorrelation analysis method^31^. The resolution at the center of the FOV in the flat-field SIM image is 101*±*3.4 nm, which is comparable to that of the Gaussian-SIM image, with a 2-fold improvement in the resolution compared to the wide field (WF) image. The resolution at the edge of the FOV is maintained at 102*±*2.2 nm with the flat-field SIM; however, the resolution drops to 108*±*1.6 nm with the Gaussian-SIM (Fig. 3g).

**Fig. 3.**
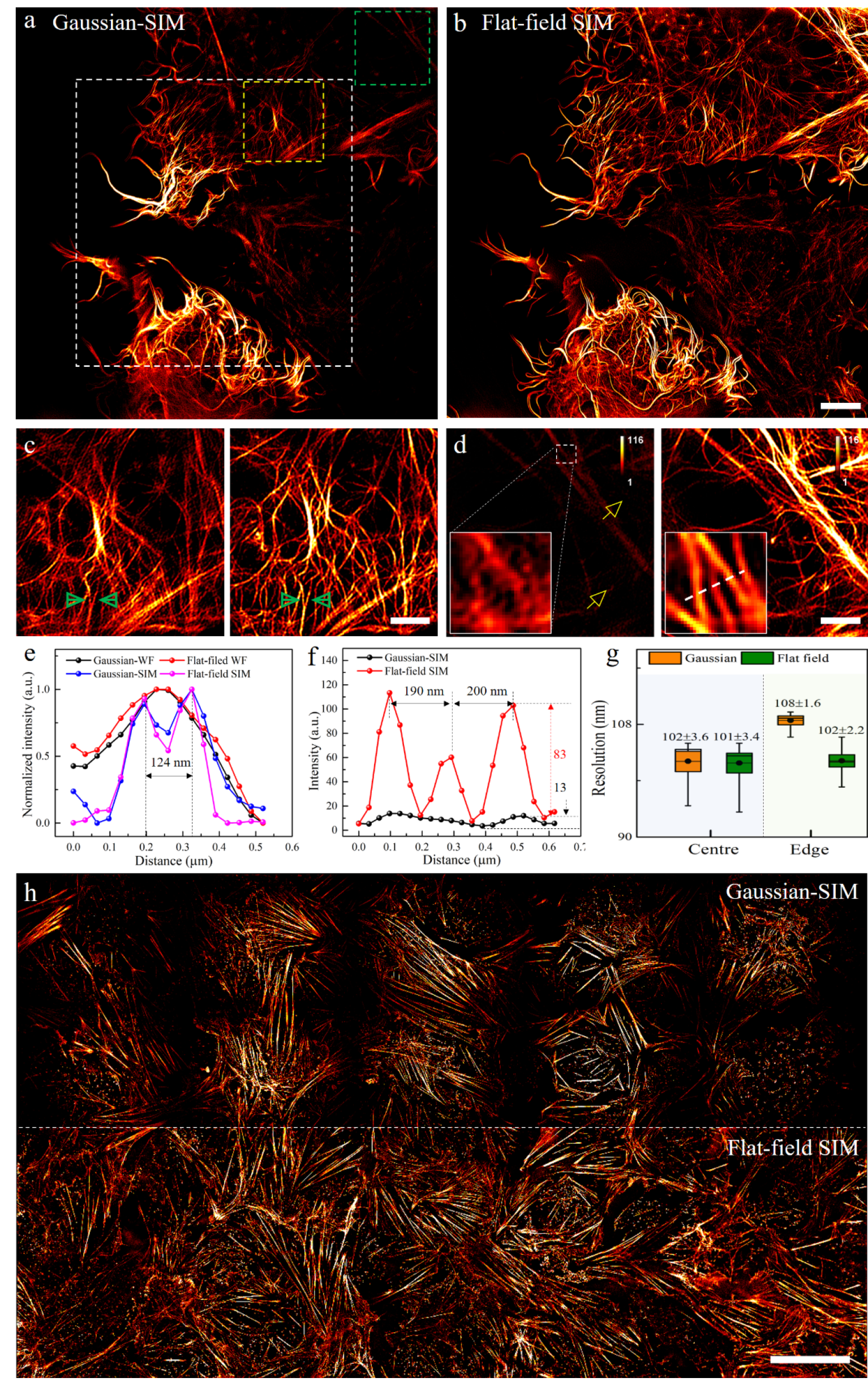
Imaging of cell microfilaments by Gassian-SIM and Flat-field SR-SIM imaging. a. The microfilaments in live U2OS cells imaged with the Gaussian-SIM show dimmer intensity and worse image quality at the edge. **b** The same region imaged with the flat-field SIM shows high-quality SR imaging at the entire FOV. The peak excitation intensities of the Gaussian-SIM and flat-field SIM are equivalent for comparison. **c** Zoomed-in views of the dotted yellow box areas in (a) by Gaussian-SIM (left) and flat-field SIM (right). **d** Zoomed-in views of the dotted green box areas at the edge in (a) by Gaussian-SIM (left) and flat-field SIM (right). **e** Normalized intensity profile along two closely lying microfilaments, indicated by the green arrows in (c). **f** Absolute intensity profile without normalization along the white-dotted line in (d). **g** Resolution obtained by decorrelation analysis at the center and edge of the field of view. **h** Multi-FOV mosaic image under the same FOV; a 5*x*5 grid of the mosaic image is shown in Fig. S10. Scale bar: 5 *μ*m in (a) and (b), 2 *μ*m in (c) and (d), 30 *μ*m in (h).

In Gaussian-SIM, a relatively uniform region in the center is usually selected to obtain an acceptable reconstruction quality (33.3*x*33 *μ*m^2^, white dotted box in Fig. 3a), ignoring the peripheral regions due to the low SNR and increased risk of artifacts. In the flat-field SIM; however, the effective FOV can be extended to 55*x*55 *μ*m^2^ due to its advantage of uniform excitation (Fig. 3b). With the flat-field SIM, the cytoskeletal microfilaments of a whole live cell were imaged for up to 15 min (Supplementary Video 1). The dynamic of microfilament growth could be observed. Fig. S9 shows a single microfilament splitting into two microfilaments. The effective FOV can be further extended with the multi-FOV stitching approach. Successful stitching requires uniform intensity to acquire images with fewer dark boundaries. As a demonstration, we acquired a 5*x*5 images with the Gaussian-SIM and flat-field SIM at the same region. As depicted in Fig. 3h and Fig. S10, the stitched Gaussian-SIM images had obvious dark boundaries at the edges, whereas the flat-field SIM enabled seamlessly stitched images with homogenous intensity and minimal image overlap.

### Whole-cell dynamics of vesicle transportation by flat-field SIM

Vesicles are carriers of both matter and signals that are vital in maintaining the exchange of material, energy, and information among cells and representing pathological conditions of live cells^32, 33^. Therefore, obtaining panoramic and dynamic images of vesicles in whole cells is important for understanding the metabolism, signal transduction, invasion, and metastasis of cancer cells. SR-SIM has been shown to be an important method in dynamic biological research due to its rapid imaging and subcellular resolution capabilities. However, the limited size of a single FOV in SR-SIM restricts its application in panoramic imaging of large whole cells. Flat-field SIM and multi-FOV stitching provide the possibility.

As shown in Fig. 4, the vesicles labeled with CD63-GFP in U2OS cells were imaged with a 3*x*1 grid of FOVs, and the white-dotted line indicates the boundary of the stitched image. To capture the dynamics and heterogeneity of the vesicles, we used time-lapse imaging of the multi-FOV. With the extended FOV and uniform excitation of flat-field SIM, we could track hundreds of vesicles in a whole cell at a temporal resolution of 1.2 s in a 52*x*110 *μ*m^2^ FOV without reducing the spatial resolution (Supplementary Video 2 and Video 3). The time-lapse image (Fig. 4a-c) shows that the vesicle distribution was nonuniform. Smaller vesicles were mainly scattered at the two ends of the cell. However, the larger vesicles were mainly distributed in the center of the cell as clusters. Furthermore, we obtained quantitative statistics based on the movement speed of three types of vesicles, namely, densely aggregated vesicles (Fig. 4d), moderately aggregated vesicles (Fig. 4e), and single vesicles (Fig. 4f). We found that the degree of vesicle aggregation is positively correlated with the movement speed, with the single vesicles showing the fastest speed. Importantly, we also observed several typical dynamic events, such as a single vesicle penetrating the cell membrane (Event 9), multiple vesicles fusing (Event 4), and single vesicles separating from aggregated vesicles (Event 13).

**Fig. 4.**
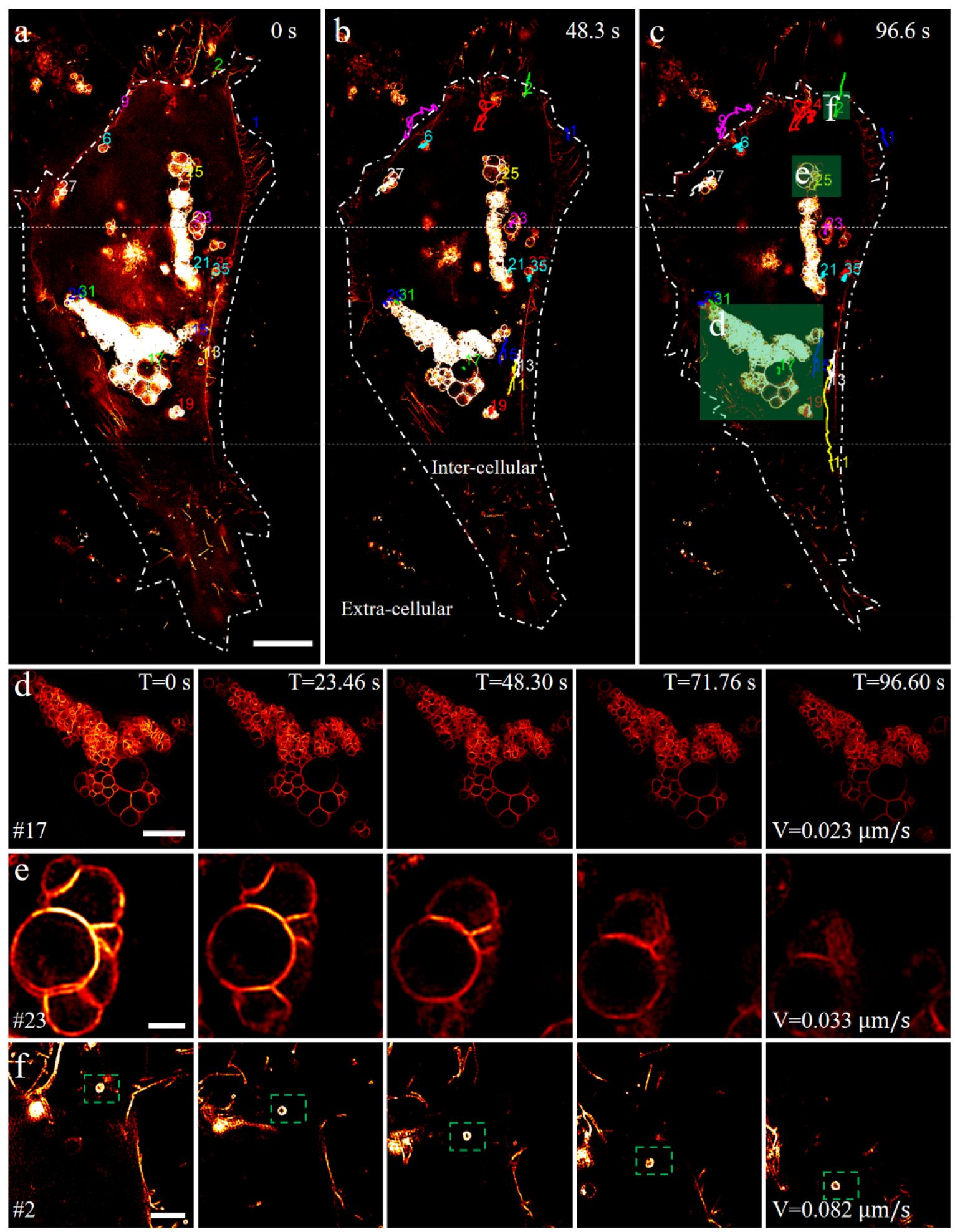
Time-lapse imaging of vesicle movement in a whole live cell by stitching three flat-field SIM images. a-c. SIM images of U2OS cells with labeled vesicles at 0 s, 48.3 s, and 96.6 s. The white dotted lines in (a)-(c) indicate the boundary of the three stitched images. **d** Zoomed-in image of the green box in (c), exhibiting the dynamic process of multiple vesicles aggregating together. **e** Breakdown of large vesicles. **f** The fast movement of a single vesicle, as indicated by the green box. Scale bar: 10 *μ*m in (a-c), 5 *μ*m in (d), 1 *μ*m in (e), 2.5 *μ*m in (f).

### Millimeter-sized SR imaging for multicell statistics and pathological diagnosis

High-throughput imaging is required to perform quantitative and unbiased statistical analysis in quantitative biology. The uniformity of the flat-field SIM images and multi-FOV stitching approach allowed the extension of the effective FOV to millimeters. It enables statistics of subcellular structures with hundreds of cells to review the cell heterogeneity. For example, mitochondria are double membrane-bound organelles that may appear as long, tubular, spherical, or elliptical structures, and the morphological abnormalities of mitochondria are essential features of tumors^34–36^.

Mitochondria structures in two cultured U2OS cell samples were imaged with flat-field SIM. One of the samples was treated with 10 μM cisplatin which is well-known to bind to DNA and critically affect the mitochondrial DNA. However, the detailed mitochondria structure changes still lack quantitative evaluation. We imaged the two samples in millimeter size covering about four hundred cells and quantitatively compared the differences in mitochondrial morphology. The stitched SR-SIM image is shown in Fig. 5a, b (additional image shown in Fig. S11). Fig. 5c, d shows dual-color images of the mitochondria and nucleus in an individual cell, which were segmented in the stitched image (segmentation method described in Supplementary Note 4). As expected, the mitochondria are distributed in a network around the nucleus and appear as thin strips with clear cristae in large numbers (Fig. 5c). However, the morphology changes significantly after treatment with cisplatin. The mitochondria exhibited fragmentation, showed low fluorescence intensity, and transformed into round, hollow structures (Fig. 5d). We quantitatively analyzed six morphological parameters, including the lengths of the minor and major axes, the ratio between the lengths, the solidity, the area, and the mean intensity. Fig. 5e-g show the distribution of the minor, major axes length and the ratio between them. The ratio was closer to 0.5 with cisplatin treatment, indicating that the mitochondria shapes were shifted from striated to rounded. The average solidity was also increased (Fig. 5h) but the area was decreased (Fig. 5i) with cisplatin treatment, indicating that the drug treatment promoted the fragmentation of the mitochondria from a multi-branched network to individual structures. Fig. 5j shows a clear difference in the mean intensities between the experimental and control groups, indicating cisplatin may impact the membrane potential^36^.

**Fig. 5.**
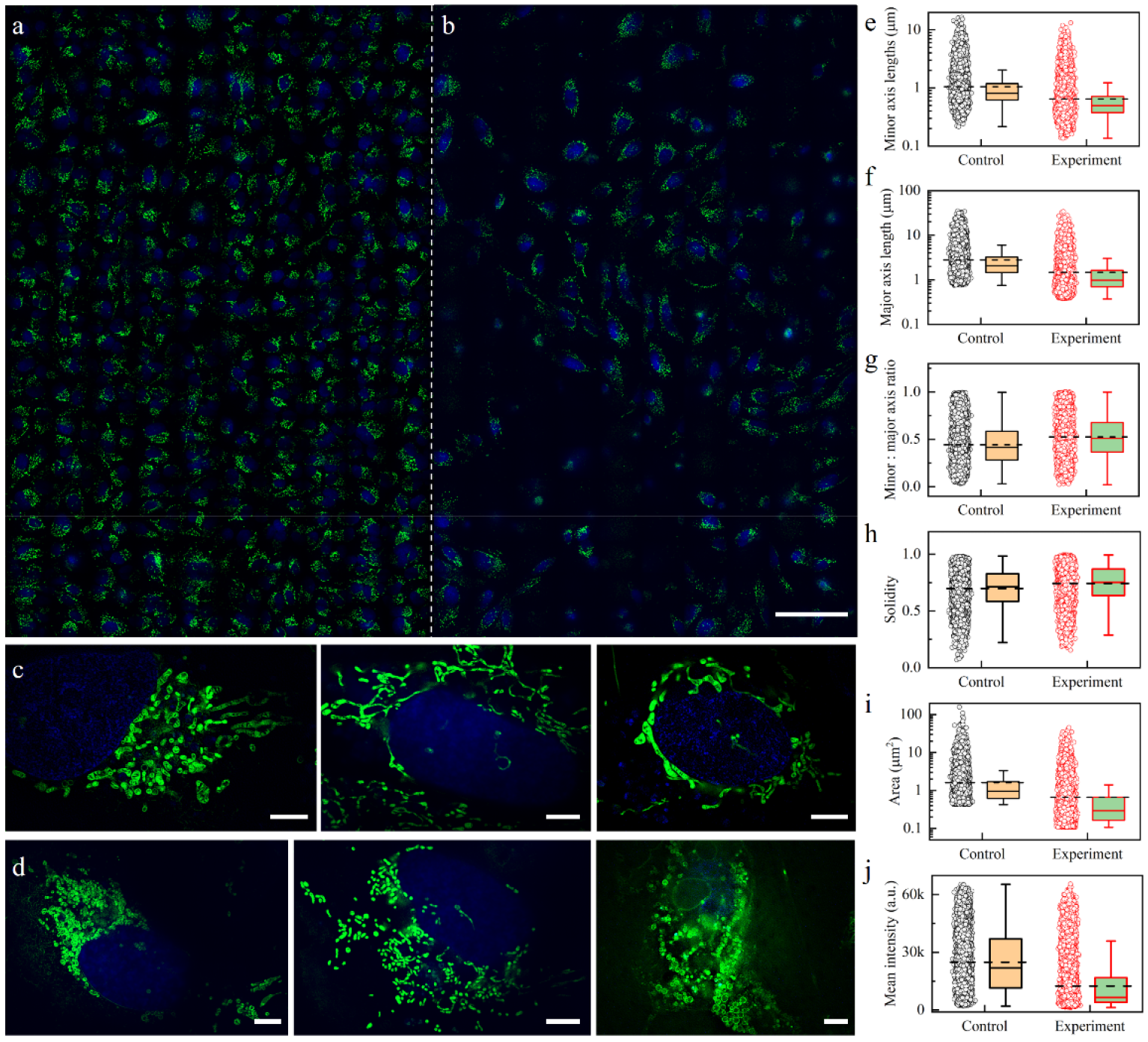
Millimeter-sized SR imaging of the cell nucleus (blue) and mitochondria (green) by stitching 20*x*20 flat-field SIM images. **a** Image of cultured U2OS cells with an FOV of 1.32*x*0.95 mm*^2^*, showing the mitochondrial structure of more than 400 cells. **b** Image of U2OS cells treated with 10 µM cisplatin as a tumor suppressor. **c** Typical cells from (a). **d** Typical cells from (b). **e-j** Distributions of minor axis length (**e**), major axis lengths (**f**), the ratios of minor axis length to major axes length (**g**), solidity (**h**), area (**i**), and mean fluorescence intensity (**j**) of the mitochondria in (a) and (b). Scale bar: 100 *μ*m in (a) and (b), 5 *μ*m in (c, d).

With the high-resolution and high-throughput imaging advantages of flat-field SIM, we conducted imaging of fluorescence in situ hybridization (FISH) pathology slides up to millimeter size. As shown in Fig. S12, two close FISH probes are indistinguishable under WF microscopy (Fig. S12b, d). In contrast, under flat-field SIM, the red and green FISH probes can be resolved into separated signals (intensity profiles shown in Fig. S12c, e). The increased spatial resolution allows for more precise enumeration and enhances the diagnostic accuracy of FISH for tumors, particularly for probes with distances smaller than the diffraction limit. We used a 10*x*10 grid of stitched images and an image mosaic approach to realize super-resolution imaging of more than 800 non-small cell lung cancer sections, providing a sufficient sample for statistical analysis (Fig. S12f).

### Selective flat-field SIM to reduce photodamage

Inspired by controlled light-exposure microscopy^37^, the photodamage can be decreased by introducing selective illumination in the region of interest. According to the properties of the joint spatial-temporal light modulation approach proposed in this paper, we can customize both the region of interest and illumination intensity. As depicted in Fig. 1d, we designed a pair of complementary illumination patterns based on the spatial distribution of the sample. For instance, by restricting the illumination region to cover only the nucleus rather than the entire FOV, potential photodamage to the mitochondria surrounding the nucleus can be reduced.

To verify the feasibility of the tunable illumination approach in reducing photodamage, we quantified the fluorescence intensity and morphology of the mitochondria over time under identical imaging conditions for a fair comparison (Fig. 6a, d). The SNR of the mitochondria in the Gaussian-SIM image rapidly decreased from an initial value of 8.8 to a value of 3.5 after 4 minutes (Fig. 6c) because of photobleaching. In contrast, the intensity of the mitochondria in the selective-SIM image gradually decreased, with the SNR decreasing from 10.2 to 8.3 (Fig. 6f). Moreover, the morphology of the mitochondria changed significantly in the Gaussian- SIM image. As shown in Fig. 6b, the mitochondria initially appeared as rod-like structures with clear cristae. Severe swelling occurred after 2 minutes; the mitochondria rounded at the ends, and the cristae disappeared, mainly attributed to phototoxicity. In contrast, the mitochondrial cristae were clearly visible after 4 minutes in the selective flat-field SIM images, and only minor background fluorescence was observed (Fig. 6e). The ratio of the lengths of the minor to major axes (suggesting the degree of phototoxicity) increased from 0.25 to 1 (75% change) in the Gaussian-SIM image, whereas the ratio changed by only 21% (from 0.31 to 0.52) in the selective flat-field SIM image due to the lower light applied to the live cells. The fluorescence intensity and morphology changes indicate that selective illumination can reduce photodamage to live cells compared with full-field illumination. The experiment was repeated five times, and the results of all experiments showed similar characteristics and performance.

**Fig. 6.**
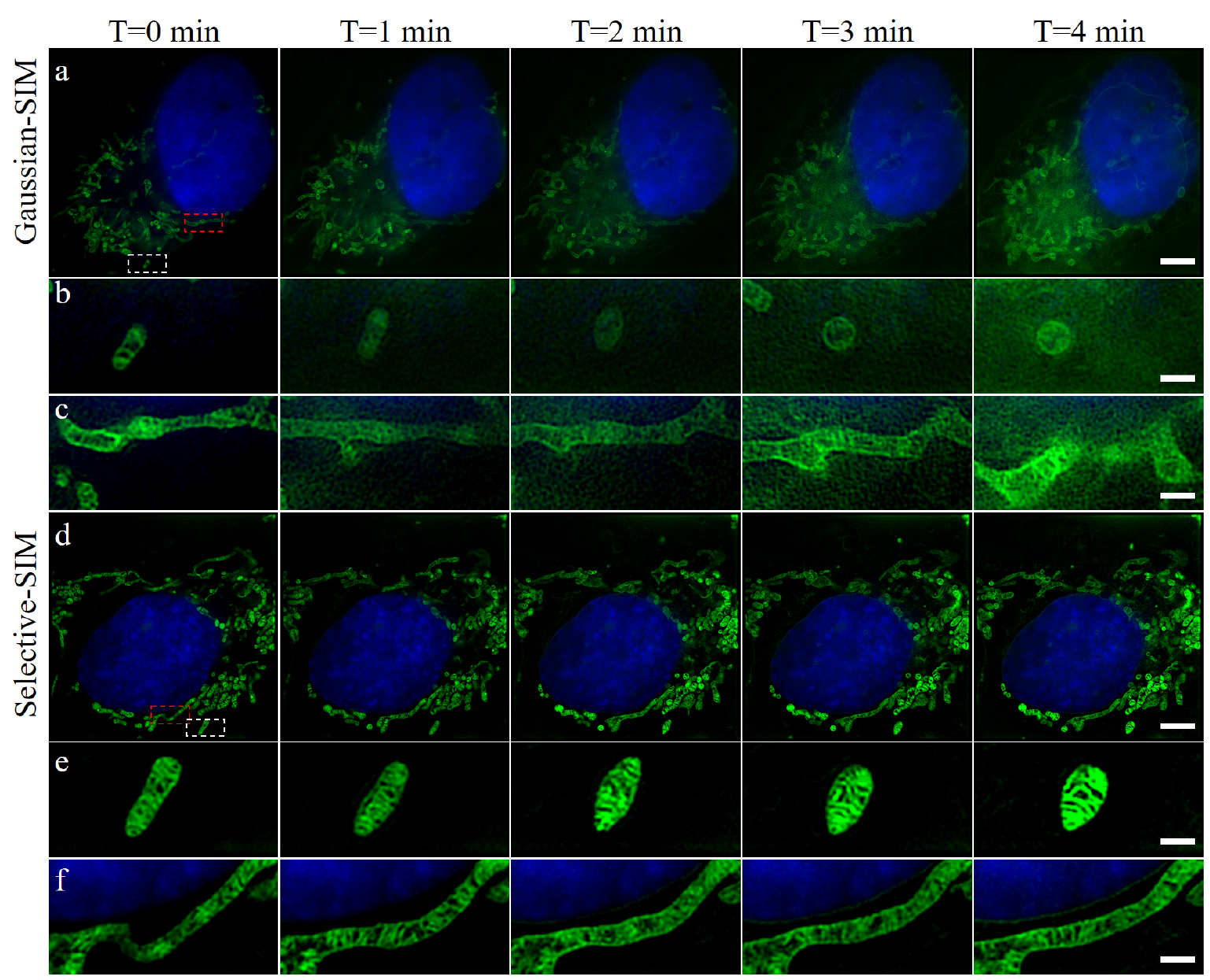
Time-lapse two-color imaging of the cell nucleus (blue) and mitochondria (green) in human osteosarcoma cells by selective flat-field SIM. a. Two-color imaging with the entire FOV illuminated by two lasers. **b, c** Zoomed-in views of the white and red boxes in the lower panels, highlighting the deformed mitochondrial morphology by photodamage. **d** Two-color imaging with the 405 nm laser illuminating only the nucleus region and the 488 nm laser illuminating the other region. **e, f** Zoomed-in views of the white and red boxes in the lower panels, showing fewer morphological changes in the mitochondria. Scale bar: 5 *μ*m in (a) and (d), 1 *μ*m in (b, c, e, f).

## Discussion

We developed flat-field SIM, a uniform and selective field excitation technique based on a joint spatial-temporal light modulation method. Compared with state-of-the-art uniform illumination schemes, the JSTLM method does not impact the wavefront of the incident light during the transformation from Gaussian to flat-field illumination, thereby ensuring high contrast in the interference fringes in the SIM images. The flat-field SIM provides homogeneous excitation across an approximately 4*x* larger FOV than the Gaussian-SIM while eliminating the position-dependent resolution and noise-related artifacts. In our flat-field SIM, the FOV is limited to 52 *x*52 *μ*m^2^ due to the limited size of the FLC-SLM (SXGA, 1280 *x* 1024 pixels, 13.6 *μ*m /pixel). By employing a larger surface FLC-SLM (ForthDD’s 4K, 2048 *x* 2048 pixels, 8.2 *μ*m/pixel), 133*x*133 *μ*m^2^ full-target coverage of the camera (2048*x*2048 pixels, 6.5 *μ*m/pixel) can be achieved with a 100x objective.

The proposed flat-field SIM approach can be easily implemented with any interferometric SIM microscope without additional optical hardware or increased costs. Although the flat-field illumination is superior to Gaussian illumination in terms of uniformity and effective FOV, the energy of the incident laser was reduced by 35% due to the inherent characteristics of the JSTLM method. To realize automatic ROI selection for field-flat SIM, the drive board of FLC-SLM can be replaced with a real-time loading function (R5, ForthDD, UK) or with a DMD for SIM. ROI illumination mode, pixel-level illumination cannot be realized due to the diffraction limitation and coherent illumination.

Benefiting from the large FOV achieved with the flat-field excitation approach, the vesicle distribution within intact live cells can be dynamically tracked over long periods. Flat-field excitation also has advantages for high-throughput super-resolution imaging with multi-FOV stitching, enabling the quantitative analysis of samples spanning multiple FOVs, such as groups of cells, pathological slides, and tissues. Combining multi-FOV stitching with flat-field SIM, super-resolution imaging of a 1.32 *x*0.95 mm^2^ FOV was achieved. The proposed multi-FOV stitching approach and flat-field SIM technology were used to quantitatively analyze mitochondrial morphology in hundreds of live osteosarcoma cells. The results showed that the mitochondrial morphology changed significantly after treatment with tumor inhibitors. The observation and statistical analysis of FISH pathological sections showed that the proposed method has higher positioning and counting accuracy than fluorescence microscopy methods, indicating its potential advantages in pathological diagnosis.

## Conclusion

In summary, the JSTLM approach is a versatile, innovative, and easy-to-implement illumination method that preserves the wavefront of the incident light during intensity modulation. With the JSTLM method, flat-field and selective illumination were achieved in SIM. Furthermore, flat-field SIM can be combined with ASOM to achieve large FOV image stitching without moving the stage to improve the throughput of SIM imaging^38^. By combining the JSTLM method with high dynamic range imaging, the dynamic range of the SIM can theoretically be increased by 40 dB^39^. With arbitrary regions of interest and the specified excitation intensity of the JSTLM method, we can combine SIM with other fluorescence interference techniques, including fluorescence recovery after photobleaching (FRAP) and fluorescence loss in photobleaching (FLIP)^40^, and develop event-driven SIM imaging methods to further reduce photobleaching and phototoxicity to live cells^41,42^.

## Methods

### Experimental setup

The custom flat-field SIM system is schematically illustrated in Fig. 1a. It is based on a commercially available inverted fluorescence microscope (IX83 or IX73, Olympus Life Science, Japan). A multicolor laser combiner (L6Cc, Oxxius, France) with four laser lines was used as the light source. In brief, four collinearly combined laser beams (200 mW) with wavelengths of 405 nm, 488 nm, 561 nm, and 637 nm were directed through an acoustic-optical tunable filter (AOTF, AA Opto-Electronic, France) for rapid switching and precise adjustment of the illumination power. A polarization-maintaining optical fiber (PM-S405XP+, Coherent, USA) was used to couple the four-color laser beams. The beam was then expanded by an achromatic objective lens (L1, TL2X-SAP, Thorlabs, USA). The expanded Gaussian beam was projected on the ferroelectric liquid crystal space light modulator (FLC-SLM, SXGA-3DM, Forth Dimension Displays, UK). The FLC-SLM is a pure-phase spatial light modulator with only 0 and *π* phases. When integrated with a polarizing beam-splitting (PBS) prism, it can function as a diffraction grating. The incident light diffracted by the grating pattern was preloaded in the FLC-SLM. The diffracted beams were focused by a Fourier lens (L2, ACA254-300-A-ML, Thorlabs, USA) onto the intermediate pupil plane, where a stop mask was placed to permit the passage of only *±*1 order beam pairs; then, the beams passed through a customized polarization rotator (PR, Union Optic, China) to maintain the polarization of the *±*1 order beam to S-polarization. The *±*1 order beam was passed through a posterior mirror set (L3, ACA254-150-A-ML, Thorlabs, USA) and a tube lens (L4, ACA254-200-A-ML, Thorlabs, USA) to focus on the back focal plane of the objective and interfere at the sample plane. The 2*x* demagnified telescope system consisted of a Fourier lens (L2) and posterior mirror set (L3), while a tube lens (L4) and 100 *x* objective lens create a 111 *x* demagnified projection, thereby illuminating the light path to achieve a 222*x* conjugate demagnified projection for the SLM. A pair of 3 mm thick polarization-preserving dichroic mirrors (ZT405/473- 488/561/640rpc-Phase, Chroma, USA) were placed orthogonally to maintain the polarization. The fluorescence images were collected by an oil-immersion objective (TIRF100*x*NA1.5 and 60*x*NA1.42, Olympus Life Science, Japan). The sample was placed on an X-Y motorized stage (H117P1XD/E, PRIOR Scientific, USA). The raw images were captured using a 2048*x*2048 pixel^2^ sCMOS camera (ORCA-Flash4.0 V2, Hamamatsu, Japan; BSI, and Teledyne Photometrics, USA). The optical pixel size was approximately 108 nm with a 60*x* objective lens and 65 nm with a 100*x* objective lens. The custom flat-field SIM microscope system is depicted in Fig. S13. The dimensions of the SIM module are 45*x*45*x*9 cm^3^.

### Fluorescent sample preparation

*Beads.* Using a pipette gun, 2 *μ*L of the fluorescent bead solution (diameter 100 nm, yellow-green fluorescent 505/515, F8803, Thermo Fisher Scientific) and 2 mL of the alcohol solution were diluted 1000-fold and placed into 5 mL centrifuge tubes. To disperse any agglomerated beads, the solution of fluorescent beads in the diluted centrifuge tube was subjected to ultrasonic treatment for 10 minutes. Then, 100 *μ*L homogeneous mixed solutions were placed into a confocal dish (#0 Glass, Cellvis) and incubated for 30 minutes to allow the fluorescent beads to adhere to the bottom of the dish. The samples were utilized for SIM imaging after adding 2 mL to 3 mL of aqueous solution.

*Rhodamine 123 fluorescein.* DMSO (1 mL) was added to rhodamine 123 (C2007, Beyotime Biotechnology, China) to prepare a stock solution, and then pure water was used to dilute the solution 1000 times to prepare a working solution. A total of 10 *μ*L working solution was added to 29 mm glass-bottomed cell culture dishes (#0 Glass, Cellvis), which were covered with coverslips (12-545-100, Thermo Fisher, USA).

### Cell culture and labeling

*Cell culture.* The U2OS osteosarcoma cell line was purchased from the Stem Cell Bank at the Chinese Academy of Sciences. The U2OS cells were cultured in McCoy’s 5A (modified) medium (Thermo Fisher, USA) supplemented with 1% penicillin G, streptomycin (Sangon, China), and 10% fetal bovine serum (Thermo Fisher, Australia) at 37 °C in a 5% (v/v) CO2 environment. The cells were transfected using Lipofectamine 8000 (Beyotime Biotechnology, China) under the manufacturer’s protocol.

*Microfilament labeling.* The cell microfilament was labeled by the LifeAct-GFP vector. After the cells reached 80% confluence, the cells were seeded at a 1:3 ratio into 29 mm glass bottom dishes (#0, Cellvis, China). The cells were cultured for 24 hours before transfection and an additional 24 hours before the experiments.

*Vesicle labeling.* The cell vesicle was labeled by the CD63-GFP vector. After the cells reached 80% confluence, the cells were seeded at a 1:3 ratio into 29 mm glass bottom dishes (#0, Cellvis, China). The cells were cultured for 24 hours before transfection and an additional 24 hours before the experiments.

*Mitochondria and nucleus labeling.* Cell mitochondria were labeled with MitoTracker Green FM (M7514, Invitrogen, USA), and the operation was carried out according to the product instructions. To detect whether cisplatin affects mitochondria, U2OS cells were treated with cisplatin at a concentration of 10 μM for 24 h, and the nuclei of the cells were labeled with NucBlue (R37605, Thermo Fisher Scientific, USA) and observed with an SIM microscope.

*FISH labeling.* Non-small cell lung cancer tissue sections were provided by the Department of Pathology, Fudan University Cancer Hospital, and the production date was 2021. The paraffin-embedded non-small cell lung cancer sections were baked and soaked for deparaffinization. After rinsing, protease digestion, buffer washing, and air drying, the EGFR/CSP7 dual-color probe (F.01009-01, Guangzhou Anbiping Pharmaceutical Technology, China) was hybridized with the section. The hybridization reaction was carried out with an in situ hybridization instrument (Abbott, USA). After hybridization, strict washing steps were performed to remove unbound FISH probes. The nuclei were stained with DAPI (C1005, Beyotime Biotechnology, China) after drying.

### Image acquisition, reconstruction, and statistical analysis

*Image acquisition.* The beads were imaged with TIRF-SIM under a 488 nm laser using a 100x oil objective configuration and an exposure time of 20 ms. Rhodamine fluorescein imaging was performed under a 2D-SIM configuration. To ensure equivalent peak power at the focal plane, the output power of the 488 nm laser was adjusted to 130 mW for Gaussian illumination and 200 mW for flat-field illumination. This results in equivalent fluorescence intensities in the center of the FOV for both Gaussian and flat-field illumination, facilitating a fair comparison. Microfilaments were also imaged at 20 ms integration times by the TIRF-SIM. The vesicles, mitochondria, and nucleus were imaged under a 2D-SIM configuration. The microfilaments, vesicles, and mitochondria were excited by a 488 nm laser, and nuclei labeled by NucBlue were excited by a 405 nm laser. The FISH imaging was conducted with a 20 ms exposure time using a 60*x* oil objective and 2D-SIM configuration. For multi-FOV image stitching, raw images were acquired with a 5% overlap between adjacent FOVs.

*Image reconstruction.* The SIM images were reconstructed by a HiFi-SIM software package^41^ developed by our group. The reconstruction parameter for ‘attStrength’ was set to 0.92 in TIRF-SIM and 0.98 in the 2D-SIM configuration, while the default values were used for the other reconstruction parameters. To enable video SIM reconstruction, we rewrote the HiFi-SIM code, and the code is available from the authors upon reasonable request.

*Image processing and analysis.* Image processing and analysis were primarily performed using FiJi. For multi-FOV image stitching, we utilized an open-source toolkit (FiJi->Plugins->Stitching->Grid/Collection stitching). The tile overlap stitching parameter was set to 5%, linear blending was selected as the fusion method, and the regression threshold was set to 0.3, while the default values were used for the other parameters. For vesicle motion trajectory analysis, the Tracking open toolkit was used (FiJi->Plugins->Tracking->Manual Tracking). The ‘centering correction’ was selected as the local maximum, the main parameter of the ‘time interval’ was set to 0.36 s, and the ‘x/y calibration’ was set to 0.325 *μ*m . Statistical analysis of mitochondrial morphology was performed with a custom Python 3.7 code, and the code is available from the authors upon reasonable request.

### Statistical analysis of the peak SNR of the fluorescent beads

To quantitatively compare the SNR distribution characteristics of the fluorescent beads under Gaussian and flat-field illumination, it is necessary to precisely determine the position of each bead and calculate its corresponding SNR value. Due to diffraction limitations and the sparse distribution of the 100 nm fluorescent beads utilized in the experiment, a single-molecule localization method was employed for data processing. Gaussian fitting was used to localize the single beads, and the fitting equation is

described as

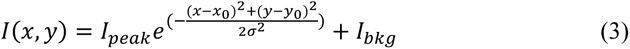

Here, the ThunderSTORM plugin (FiJi->Plugins->ThunderSTORM) was utilized to extract the main parameters, including the fluorescent bead location (*x_0_*, *y_0_*), the width of the Gaussian kernel *σ*, the peak intensity of the signal *I_peak_*, the intensity of the background signal *𝛼_bkg_*, and the standard deviation of the background signal *𝛼_bkg_*. These parameters were used to calculate the peak signal-to-noise ratio of each fluorescent bead across the full field of view, which was used in the quantitative analysis. The peak signal-to-noise ratio can be calculated as

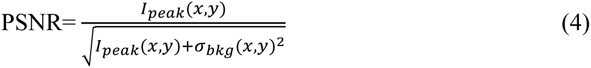

where *𝛼_bkg_* is the standard deviation of the background signal, which is calculated based on the surrounding pixels near the center of the fluorescent beads. In the ThunderSTORM plugin, Gaussian is used as the localization method, and the least squares fitting method is selected. The localization results of the fluorescent beads under both Gaussian and flat-field illumination are shown in Fig. S8.

## Data Availability

All the data that support the findings of this study are available from the corresponding author upon reasonable request.

## Code Availability

All code supporting the findings of this study is available from the corresponding authors upon reasonable request.

## Acknowledgments

The National Natural Science Foundation of China [grant no. 62205367, 62141506], Suzhou Basic Research Pilot Project [grant no. SSD2023006], and National Key Research and Development Program of China [grant no. 2017YFC0110100].

## Notes

### Competing Interest Statement

The authors have declared no competing interest.

